# Amorphous silicon resistors enable smaller pixels in photovoltaic retinal prosthesis

**DOI:** 10.1101/2025.05.01.651774

**Authors:** Andrew Shin, Nathan Jensen, Emma Butt, Jeonghyun An, Davis Pham-Howard, Ludwig Galambos, Keith Mathieson, Theodore Kamins, Daniel Palanker

**Affiliations:** Department of Materials Science and Engineering, Stanford University, Stanford, CA, US; Department of Electrical Engineering, Stanford University, Stanford, CA, US; Institute of Photonics, Department of Physics, University of Strathclyde, Glasgow, UK; Hansen Experimental Physics Laboratory, Stanford University, Stanford, CA, US; Department of Ophthalmology, Stanford University, Stanford, CA, US

## Abstract

**Objective:** Clinical trials of the photovoltaic subretinal prosthesis PRIMA demonstrated feasibility of prosthetic central vision with resolution matching its 100µm pixel size. To improve prosthetic acuity further, pixel size should be decreased. However, there are multiple challenges, one of which is related to accommodating a compact shunt resistor within each pixel that discharges the electrodes between stimulation pulses and helps increase the contrast of the electric field pattern. Unfortunately, standard materials used in integrated circuit resistors do not match the resistivity required for small photovoltaic pixels. Therefore, we used a novel material – doped amorphous silicon (a-Si) and integrated it into photovoltaic arrays with pixel sizes down to 20µm.

**Approach:** To fit within a few µm^2^ area of the pixels and provide resistance in the MΩ range, the material should have sheet resistance of a few hundred kΩ/sq, which translates to resistivity of a few Ω*cm. The a-Si layer was deposited by low-pressure chemical vapor deposition (LPCVD) and its resistivity was adjusted by PH_3_ doping before encapsulating the resistors between SiO_2_ and SiC for stability in-vivo.

**Main Results:** High-resolution retinal implants with integrated shunt resistors were fabricated with values ranging from 0.75 to 4 MΩ on top of the photovoltaic pixels of 55, 40, 30 and 20 µm in size. Photoresponsivity with all pixel sizes was approximately 0.53 A/W, as high as in the arrays with no shunt resistor. The shunts shortened electrodes discharge time, with the average electric potential in electrolyte decreasing by only 21–31% when repetition rate increased from 2 to 30 Hz, as opposed to a 54–55% decrease without a shunt. Similarly, contrast of a Landolt C pattern increased from 16-22% with no shunt to 22–34% with a shunt. Further improvement in contrast is expected with pillar electrodes and local returns within each pixel.

**Significance:** Miniature shunt resistors in a MOhm range can be fabricated from doped a-Si in a process compatible with manufacturing of photovoltaic arrays. The shunt resistors improved current injection and spatial contrast at video frame rates, without compromising the photoresponsivity. These advances are critical for scaling pixel sizes below 100 µm to improve visual acuity of prosthetic vision.

## Introduction

Retinal degenerative diseases, such as Retinitis Pigmentosa and Age-related Macular Degeneration (AMD) are among the leading causes of incurable blindness today [1]. They result in a progressive loss of photoreceptors, but the inner retinal neurons largely survive, preserving many aspects of natural retinal signal processing [2-6]. Retinal prostheses are designed to reintroduce visual information into the retina by electrical stimulation of these remaining neurons. The photovoltaic subretinal prosthesis was developed as a substitute for the lost photoreceptors to restore central vision in the geographic atrophy (GA) region of AMD patients. Each pixel in this implant converts light into electric current to stimulate the second-order retinal neurons, primarily the bipolar cells (Figure 1B) [7]. Photodiodes require much brighter light than ambient for neural stimulation, and electric current needs to be pulsed to preserve the charge balance at the electrode/electrolyte interface. Therefore, images captured by a camera are projected from augmented-reality (AR) glasses onto the implant using pulsed intensified light (Figure 1A). To avoid any confounding effects of this intense light on remaining photoreceptors outside the GA, its wavelength is in the near-infrared part of the spectrum (880 nm). Since the photovoltaic implant is completely wireless, its implantation is quite simple, and it does not affect the surrounding residual photoreceptors [8,9].

**Figure 1.**
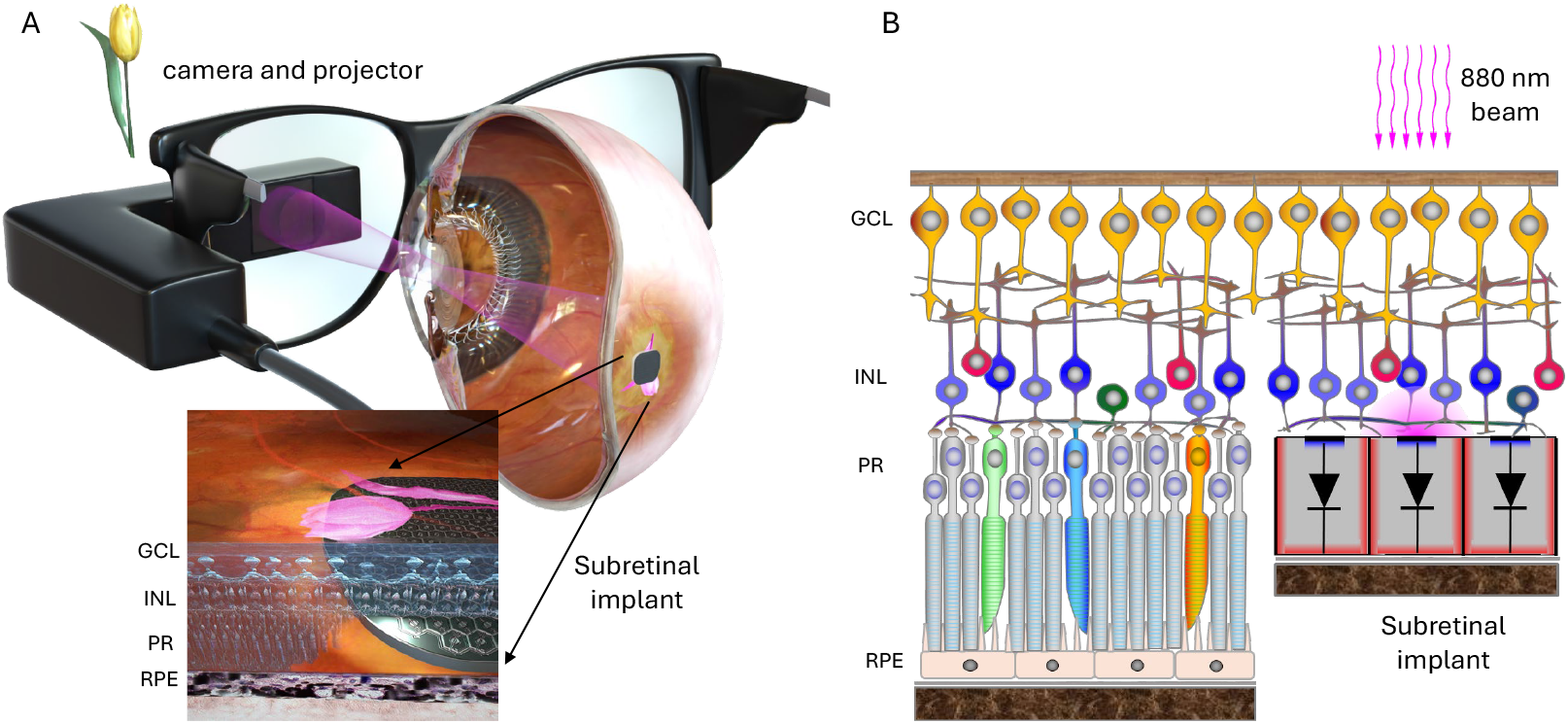
System diagram of the photovoltaic retinal prosthesis. (A) Camera captures the visual scene and processed images are projected onto the subretinal implant using a pulsed near-infrared (880 nm) beam. (B) Photovoltaic pixels convert incoming light into electrical current flowing through the retina between the active and return electrode. Resulting electric field polarizes the neurons in the Inner Nuclear Layer (INL) of the retina.

In a series of two clinical trials, this photovoltaic implant (PRIMA, Pixium Vision) was implanted in 43 AMD patients suffering from central vision loss due to GA. Prosthetic vision enabled patients to reliably recognize letters and words, with a visual acuity of 20/420, which closely matches the sampling limit of spatial resolution enabled by its 100 µm pixels [8-10]. While these landmark results represent an important proof of concept for prosthetic vision, the number of AMD patients who could benefit from this technology would grow significantly if the implant resolution was increased. For example, to provide an acuity of 20/100, pixels should be no larger than 25 µm.

Unfortunately, smaller pixels with a planar photodiode structure exhibit faster recombination of photogenerated charge carriers on the pixel walls, leading to reduced photoresponsivity. To overcome this limitation, we developed doping in vertical trenches separating the photodiodes, which resulted in 0.51 A/W photoresponsivity for all pixel sizes, down to 20 µm (an improvement from 0.25 A/W) [11]. To adapt such arrays for a retinal prosthesis operating at 30 Hz frame rate, a shunt resistor should be added to discharge the electrode capacitance between the pulses [12] and also to provide a pathway for return current via dark pixels. However, fitting a MOhm-range resistor in a small area of the pixel requires materials with sheet resistance outside the conventional range. Here, we describe the development of novel amorphous silicon-based (a-Si) shunt resistors and their integration with photodiodes in each pixel of the photovoltaic array.

## Methods

### Optimization of the shunt resistor

Clinically, photovoltaic implants operate at 30 Hz, with 1-10 ms pulses at 3.5 mW/mm^2^ irradiance [8,9]. Under these conditions, without a shunt resistor, charge accumulates at the electrode-electrolyte interface and the voltage across the photodiode builds up, making it conductive enough to drain the excess current and maintain the charge balance during the cycle (Figure 2A-C). This reduces the charge injection into the retina and thereby lowers the efficiency of neural stimulation (Figure 2C). To overcome this limitation, the shunt resistor is connected in parallel to the photodiode, allowing a discharge of the capacitive interface between the stimulating electrode and electrolyte (tissue), as shown in Figure 2 C-E. Since in this configuration the shunt resistors provide an alternative path for electric current, the optimum value of its resistance should be selected to maximize the charge injection into the retina, as described previously [13]. For pixels of 20, 30, 40 and 55 µm in width, the optimum resistances were found to be 4.0, 1.7, 1.0 and 0.75 MΩ, respectively. As can be seen in Supplementary Figure 1, deviations from these values within +/-30% during fabrication should not significantly affect the device performance.

**Figure 2.**
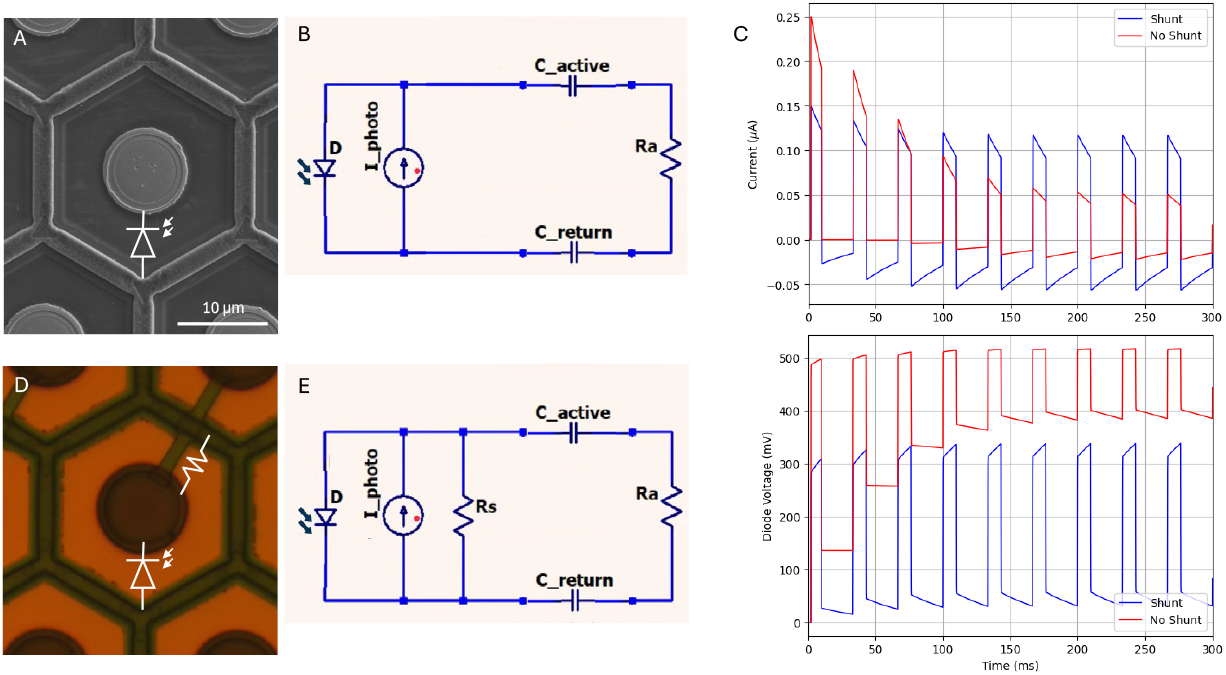
Modeled performance of the photodiode pixels with and without a shunt resistor. (A) Array of 30 µm pixels with photodiodes without shunt resistors. (B) Equivalent circuit diagram of the pixel–tissue interface, incorporating the electrode–electrolyte capacitance. (C) Electric current and voltage across the diode generated by pulses of light at 1.0 mW/mm^2^ irradiance and 30 Hz repetition rate. (D) Array of 30 µm pixels with shunt resistors connecting the active and return electrodes. (E) The corresponding circuit diagram of the implant–tissue interface including the electrode capacitance and the shunt resistor.

Another advantage of a shunt resistor in each pixel of monopolar arrays is that it provides an additional path for current to return through dark pixels, which helps improve the contrast of the electric patterns, as illustrated in Figure 3 [14]. The dark pixel in the gap of Landolt C in Figure 3C conducts current to a common return electrode, thereby reducing the electric potential in that area, compared to the non- conductive pixel. As a result, electric potential above the conductive pixel is reduced, improving the contrast of the Landolt C pattern, as shown in Figure 3D.

**Figure 3.**
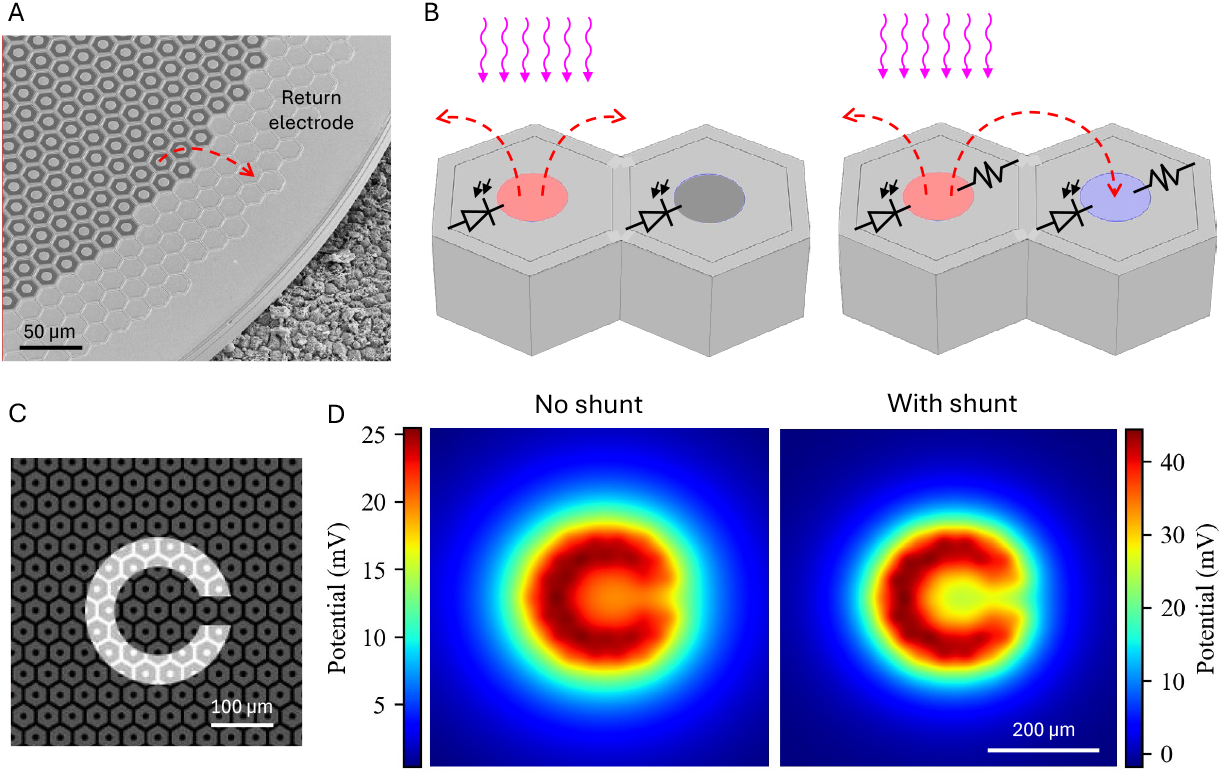
Modeled performance of photovoltaic arrays. (A) SEM image of a photodiode array with 20 µm monopolar pixels having a common peripheral return electrode. Local return electrodes within each pixel are insulated by a SiC layer. (B) Integration of the shunt resistors enables active electrodes on adjacent non-illuminated pixels act as current sinks, connected to the return electrode mesh via the shunt. (C) A Landolt-C pattern with 48 µm gap, projected onto array of 40 µm pixels. (D) Modeled electric potential in electrolyte 20 µm above the devices with and without a shunt resistor at 1.64 mW/mm^2^ irradiance and 30 Hz frame rate.

### Amorphous silicon as a new material for the shunt resistor in retinal implants

In commercial PRIMA implants, the shunt resistors in the 100 µm pixels are made from polysilicon with a resistivity in the range of 10^−3^-10^−2^ Ohm*cm (Figure 4A). However, for smaller pixels (20-40µm in size), the required shunt resistor value is much higher, and the available area is much smaller. Therefore, materials with much higher sheet resistance are required – in the range of MOhm/sq. To be integrated into the arrays, the resistor thickness should be in the 0.1-1µm range, limiting the resistivity for a MOhm resistor material to a few Ohm*cm (Figure 4A). Although single-crystal silicon can get close to this resistivity, an isolated resistor of single crystal cannot be practically formed. Metal oxides have higher resistivities (10^3^-10^8^ Ohm*cm) and hence would require much thicker films than can be accommodated within the device. Polycrystalline silicon is often used as a resistive material, but in the required range, its resistivity is difficult to control - it changes very rapidly with the dopant concentration (Figure 4A) [15]. In addition, its resistivity also varies with the layer thickness below 250 nm due to grain boundaries, which affects the reproducibility (Figure 4B) [16]. Therefore, an alternative material, compatible with silicon integrated-circuit fabrication is needed.

**Figure 4.**
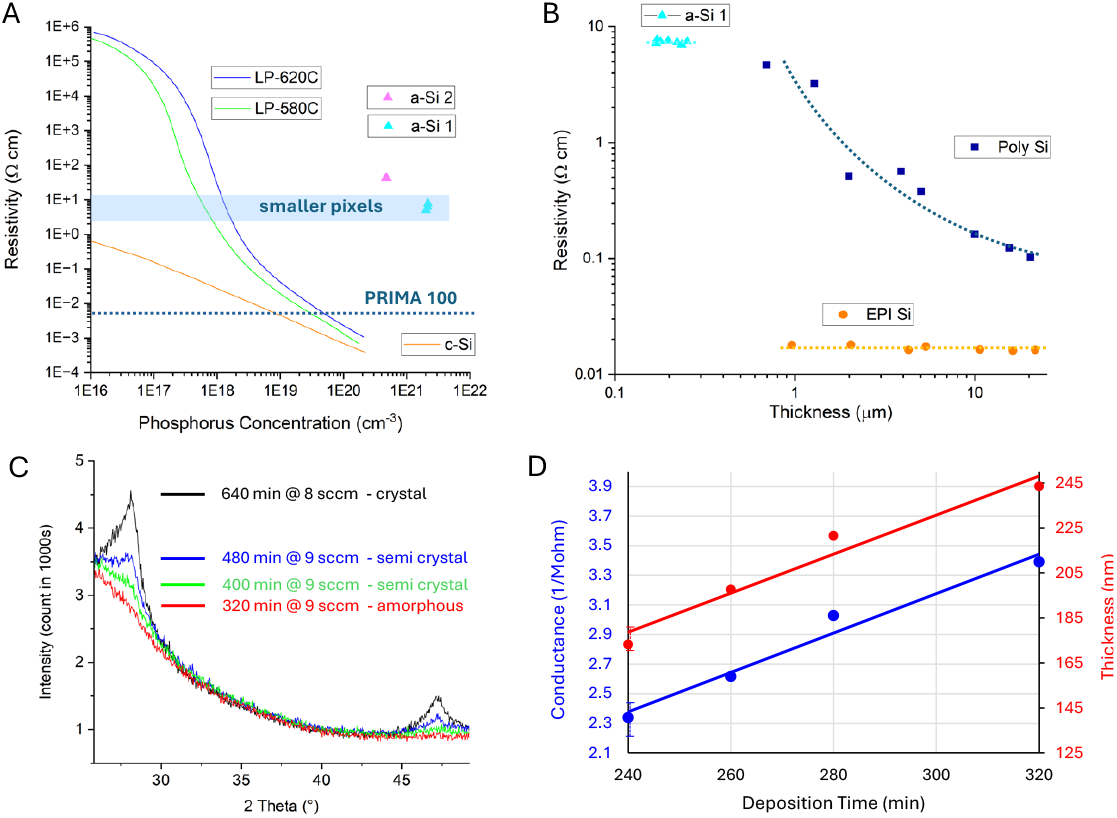
Characteristics of the resistive materials. (A) Resistivity of the phosphorus-doped single-crystalline silicon (c-Si), polycrystalline silicon, and amorphous silicon (LPCVD at 550 °C, 3 SCCM and 9 SCCM, denoted a-Si 1 and 2). For comparison, we also show the polysilicon used in the PRIMA system (100 µm pixels) and the resistivity range required for 20-50 µm pixels. (B) Variation of resistivity with the layer thickness in crystalline silicon (EPI Si), polysilicon (Poly Si), and amorphous silicon (a-Si), replotted from Fig. 5.9 in [16]. (C) X-ray Diffraction (XRD) analysis of amorphous silicon layers deposited at 550 °C with various durations. Crystalline features begin to appear beyond 400 minutes of deposition. (D) Conductance and thickness of a-Si films deposited at 9 SCCM PH_3_ flow rate. Linear scaling of conductance with thickness confirms the constant layer resistivity.

The required range of resistivity can be achieved with moderately- doped amorphous silicon, utilizing its low carrier mobility in the disordered layer. Amorphous silicon delivered by plasma-enhanced chemical vapor deposition (PECVD) is widely used as the channel of thin-film transistors in large-area electronics, such as liquid-crystal displays. However, this form of amorphous silicon contains large amount of hydrogen, which is unstable above 350 °C, making it incompatible with the 450 °C required for annealing of the metal/alloy to form stable contacts for photodiodes and resistors.

Amorphous silicon deposited by low-pressure CVD (LPCVD), on the other hand, contains very little hydrogen and is stable even within the 500 °C range. The deposition rate increases markedly with increasing temperature, but above 550 °C, the layer starts crystallizing during the time required for depositing films of a reasonable thickness (∼100 nm). With these constraints, 550 °C was chosen as the optimal deposition temperature. To determine the maximum deposition time which still avoids significant crystallization, we used X-ray diffraction (XRD). Figure 4C shows that a mixture of crystalline peaks characteristic of a polycrystalline silicon appears at 400 minutes and becomes very pronounced at 640 minutes. We used a deposition time of 320 minutes to ensure that the film is still fully amorphous (within the resolution of the x-ray system).

To adjust resistivity, layers can be doped with phosphorus or boron during deposition. Adding the dopants after deposition by ion implantation or gas-phase doping is not practical because the high temperature required for its activation would cause the amorphous layer to crystallize. To determine the dopant concentration, we used secondary ion mass spectrometry (SIMS). The desired resistivity of the amorphous silicon layer (Figure 4A) was achieved in a reproducible fashion using a PH_3_ flow rate of 1-10 SCCM (15 % PH_3_ in SiH_4_). Within the 240- 320 minutes of deposition in the LPCVD system, the layer conductance increased linearly with its thickness, indicating constant resistivity (Figure 4D).

Amorphous silicon is nearly transparent in the 800-900 nm wavelength range [17], which allows minimizing any loss of photosensitive area. However, it may be difficult to control reflection, given the multilayer antireflection stack of dielectrics is optimized for the crystalline silicon areas.

### Fabrication of the shunt resistor and its integration with photovoltaic pixels

Silicon on Insulator (SOI) wafers with 30 µm device layer and 0.5 µm Buried Oxide (BOX) layer were used for the 100 mm-sized fabrication process control in the Stanford Nanofabrication Facility (SNF). Figure 5 illustrates the fabrication process with ten mask layers, including etching and doping of the high aspect ratio vertical junction (A,B), doping of the active and return ohmic contacts (D,E), coating the anti-reflection layers and metal electrodes (F), deposition and etching the LPCVD-doped amorphous silicon (G), and releasing the devices (J,L). The fabrication process flow of the vertical p-n junction is discussed in our earlier publication [11]. The amorphous-silicon resistor layer was designed to be insulated between a 50 nm thick SiO_2_ layer from below and a 230 nm thick a-SiC layer from above (Figure 5I). Thicknesses of the SiO_2_-SiC layers was optimized to minimize the reflection of 880 nm light from the device layer (stack of 30 µm thick silicon, 0.5 µm thick buried oxide, and 0.2 µm thick titanium/platinum layer on back side).

**Figure 5.**
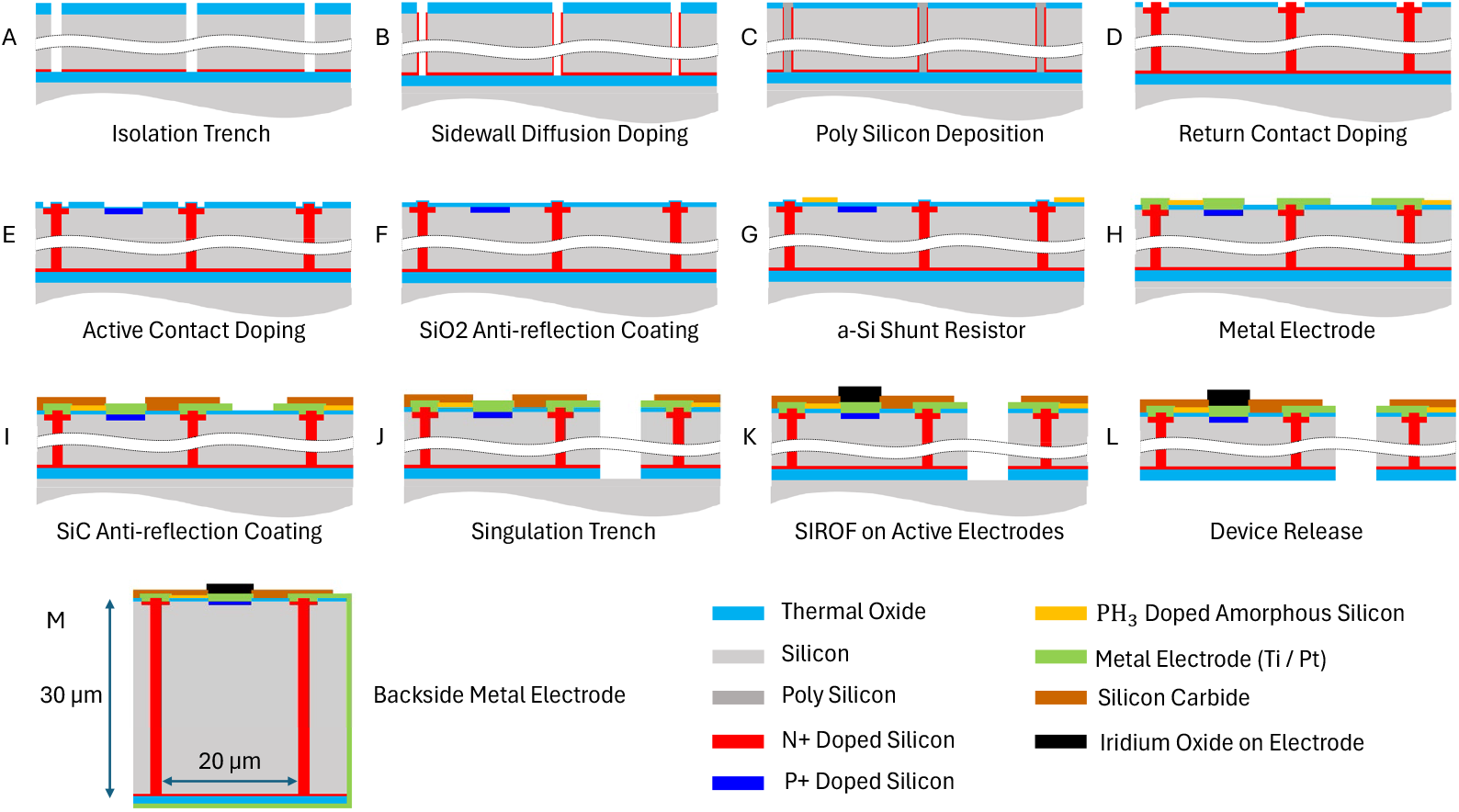
Process flow for fabrication of photodiode arrays, including the vertical p–n junctions (A-B), an a-Si shunt resistor (G), SIROF (Sputtered Iridium Oxide Film) electrodes (K) and Ti/Pt coatings on the back and side walls following the device release (M). Layer thicknesses are shown not to scale for better visibility.

Due to its limited thermal stability, the a-Si shunt resistor was added late in the front-end fabrication process – just before the metal interconnections were added (Figure 5G). After this step, the only significant thermal steps were a 450 °C forming-gas alloy/anneal and PECVD amorphous SiC deposition at 350 °C (at EIC Inc, Boston). Supplementary Figure 2 shows the changes in resistance after each of these steps. The decrease in resistance during a-SiC deposition is attributed to the limited amount of hydrogen incorporated into a-Si. These changes were reproducible and within the tolerance range of the resistor.

The sheet resistance and contact resistance after patterning, after annealing at 450 °C, and after the deposition of a-SiC were measured using Kelvin and van der Pauw test structures. To achieve the desired resistances within pixel sizes of 20 to 55 µm, we produced a-Si layers with resistance of 0.30 MOhm/square. The contact area was optimized to keep the contact resistance below 10% of the shunt resistance for each pixel size.

Devices were released from the SOI wafer after the 30 µm deep singulation trenches were defined with deep reactive ion etching (Figure 5J) and the back side of the wafer was ground with mechanical polishing and XeF_2_ etching (Figure 5L). After the release, the back side of the device was connected to the peripheral return electrode on the front side by sputtering a 200 nm thick Ti/Pt layer on the side and back of the released device (Figure 5M). For in-vivo applications, the return electrode on the back side is also coated with a 200 nm layer of sputtered iridium oxide film (SIROF) to provide large capacitance of the return electrode.

## Results

Fabricated photovoltaic arrays with shunt resistors are shown in Figure 6. Given the constant resistivity of the a-Si layer for all the devices on the wafer (0.30 MOhm/sq), the width and length of the shunt was designed to provide the required resistance for each pixel size: 0.5, 0.9, 1.8 and 3.9 MOhm for pixels of 55, 40, 30 and 20 µm, values optimized in our modeling study [21].

**Figure 6.**
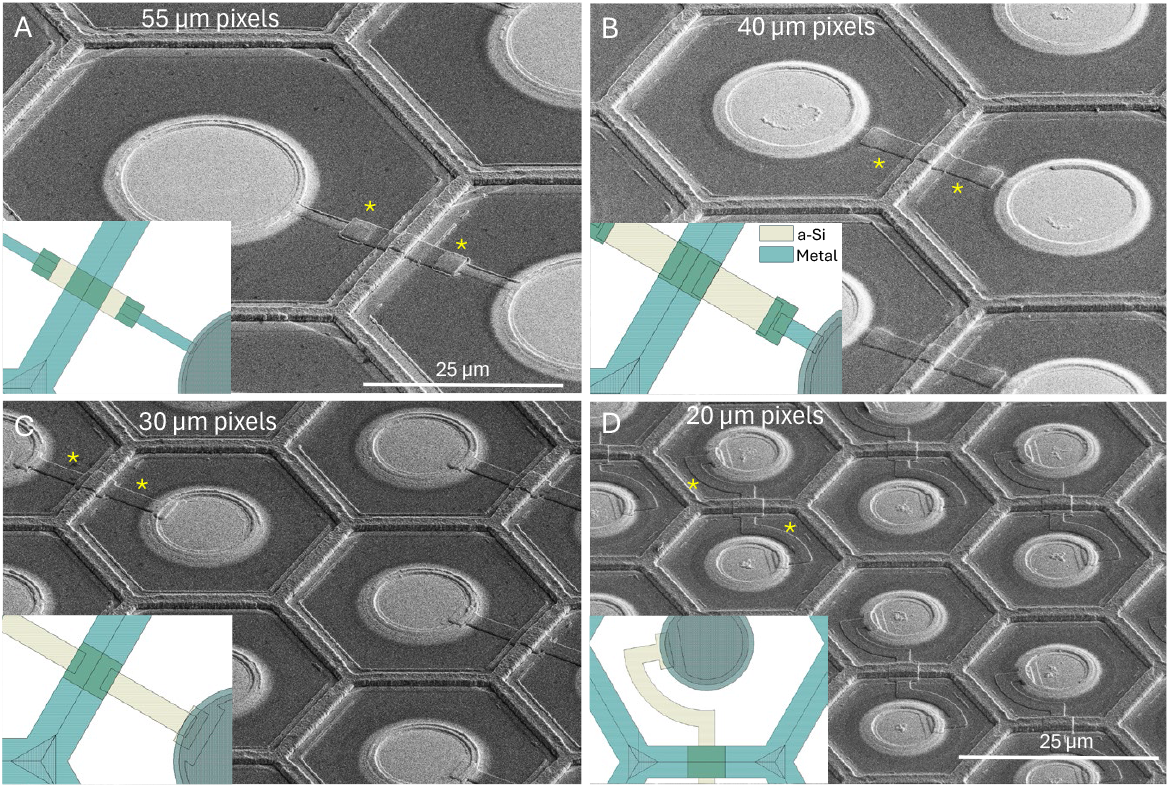
SEM images of the released devices with pixel sizes of 55 µm (A), 40 µm (B), 30 µm (C) and 20 µm (D). Shunt resistors are indicated by yellow asterisks. Inserts illustrate layout of the a-Si and metal layers for each pixel size.

### Electrical characteristics of the photodiode and shunt resistor

The I-V curves of the single-pixel test structures with and without the shunt resistor were measured from -2 to 2 V at various irradiance levels of 880 nm light – see Figure 7 for 55 µm pixels. In the dark, the photodiode test structure fabricated earlier [11] without a shunt resistor exhibited pA range current (Figure 7A), while the photodiode test structure with a shunt resistor fabricated in this work demonstrated µA range current at -2 V, as expected with the shunt resistor of 0.6 MOhm connected between the active and return electrodes (Figure 7B). When the metal connection of the shunt resistor was cut by a Focused Ion Beam (FIB), the dark current decreased to nA rather than pA range, indicating some leakage pathway on the test structure (Figure 7C). Even if the extra leakage pathway is present in the device pixels and not just in the test structures, nA leakage current is negligible compared to µA photocurrent under the laser illumination, so it should not affect the performance of the photovoltaic pixels.

**Figure 7.**
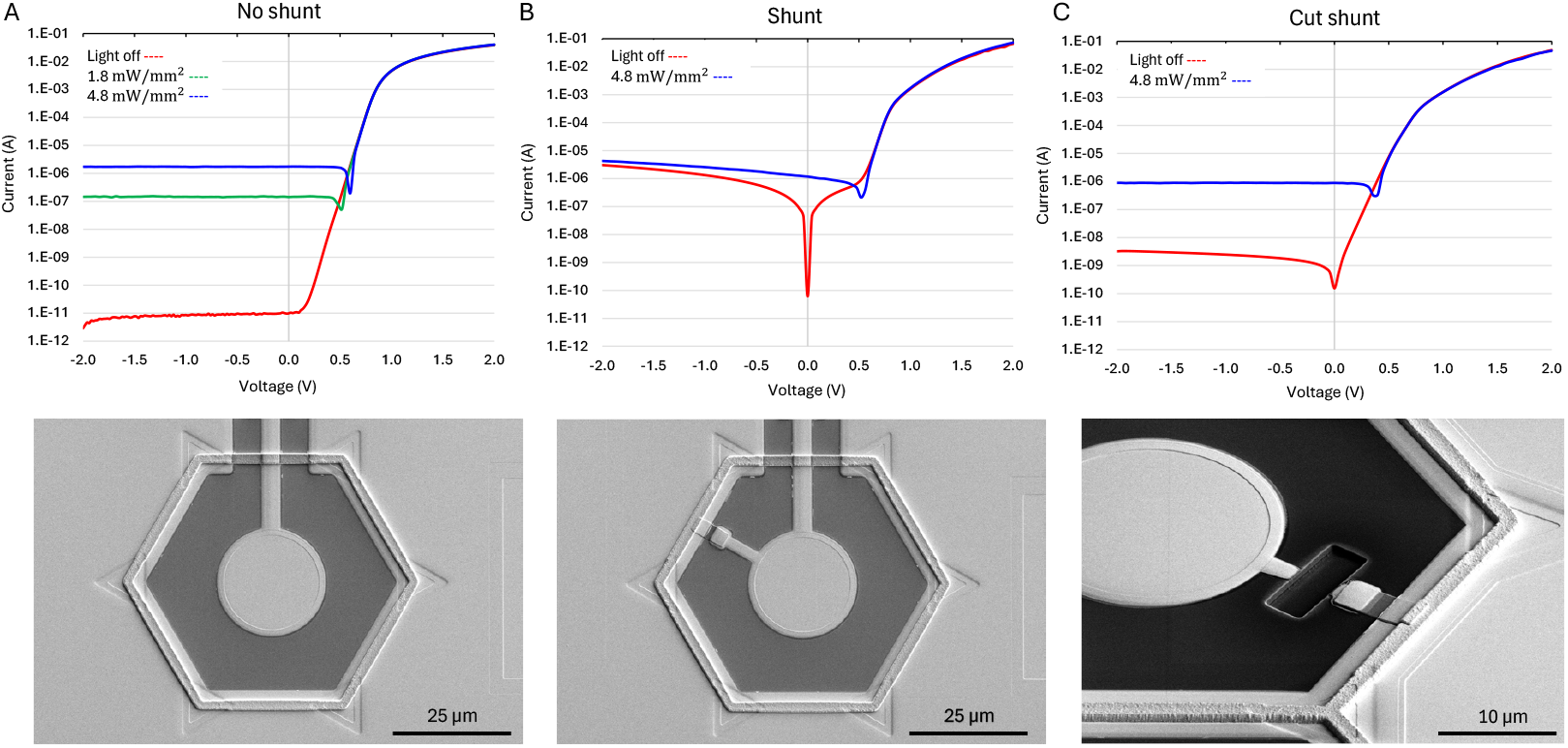
Current–voltage (IV) characteristics of the photodiodes in the dark and under 880 nm laser illumination. (A) Photodiode without a shunt resistor [11], displaying a typical forward and reverse bias behavior, including turn-on voltage and dark current. (B) Photodiode with a shunt resistor in this work, where the dark current closely matches the expected value of the shunt resistor. (C) Disconnection of the shunt resistor by a focused ion beam (FIB) reveals the residual leakage current in nA range.

This leakage current could be caused by some differences in processing to accommodate incorporation of the shunt resistor (e.g. omitting the 30 minutes N_2_ annealing process at 1000 °C after growing the thin oxide film or adding ion bombardment during patterning of the amorphous silicon) or by the FIB cutting process itself. Based on Sentaurus TCAD modeling, omitting the 30 minutes N_2_ annealing process did not significantly affect the dopant profile under the thin oxide layer. The ion bombardment, necessary to remove the native oxide before patterning the amorphous silicon and during the approximately 15% over-etch (to compensate for nonuniformities across the wafer) could result in leakage. Resistance measurements on test structures with and without a shunt resistor on 55µm pixels demonstrated that this leakage pathway did not exceed 10% of the current flowing through the shunt resistor. Residual nA-range leakage current after cutting the shunt resistor could be caused by stray metal from the FIB itself.

### Photoresponsivity of the photodiodes

To assess photoresponsivity of the photodiodes, the released devices were illuminated with 880 nm light at irradiance ranging from 0.65 to 5 mW/mm^2^. The current flowing between the active and return electrodes was measured using picoammeter (Keithley 2400). The photoresponsivity was calculated by dividing the measured current by the photosensitive area of the pixel, listed in the Table S1. At 880 nm wavelength, amorphous silicon was assumed to be transparent, so only the metal-covered areas in the pixels were assumed to block the 880 nm light.

Results are summarized in Table 1. Pixels with the shunt resistor exhibited photoresponsivity of about 0.53 A/W, like the structures without a shunt. It is important to note that to enhance the light absorption in the 30 µm-thick device, the back side of the implant was metalized with a 20 nm titanium and 200 nm platinum layer, which reflects unabsorbed light back into the device. In these measurements, MOhm-range shunt resistors did not drain any noticeable current compared to the low-resistance path of the ammeter, but the shunt did reduce the injected current in electrolyte, as described below.

**Table 1.** Average photoresponsivity of the photodiodes under the irradiance ranging from 0.6 to 4.5 mW/mm2. (A) A control implant with no shunt (N2 annealing but without a-Si deposition). (B) Array with a shunt cut by a focused ion beam. (C) Pixels with a shunt resistor.

| Pixel Size ($\mu\text{m}$) | No a-Si LPCVD, no shunt (A/W) | a-Si LPCVD, cut shunt (A/W) | a-Si LPCVD, shunt (A/W) |
| --- | --- | --- | --- |
| 55 | 0.53 ± 0.04 | 0.52 ± 0.05 | 0.53 ± 0.08 |
| 40 | 0.53 ± 0.03 | 0.54 ± 0.07 | 0.53 ± 0.05 |
| 30 | 0.53 ± 0.03 | 0.51 ± 0.06 | 0.53 ± 0.07 |
| 20 | | 0.57 ± 0.02 | 0.50 ± 0.03 |

### Charge injection into electrolyte

To evaluate the charge injection of the photovoltaic arrays into the electrolyte, the released devices were placed into diluted saline with resistivity of 640 Ohm*cm, mimicking the retinal resistivity [18]. A Landolt C pattern of light was projected onto the arrays with a gap width of 48 µm, similar to the pattern shown in Figure 3C. Patterns were projected with 9.8 ms pulses at 2 and 30 Hz at 1.64 mW/mm^2^ irradiance, and the electric potential was measured 20 µm above the device using a pipette electrode with a 5 µm opening. Electric potentials above the bright and dark parts of the C are summarized in Table 2, and the waveforms for 40 µm pixels are shown in Figure 8. As expected, for all pixel sizes, the shunt resistor increased the discharge rate of the electrodes between the pulses, which reduced the voltage across the diode and thereby made the electric current and potential in the medium more square during the pulse (Figure 8). With a shunt, the mean of current during the pulse at 30 Hz dropped by 21-31% from that at 2 Hz. Without a shunt, the waveforms exhibit a steeper drop during the pulse due to opening of the diode upon accumulation of charge on the electrodes. This is also associated with a bigger decline when transitioning to higher repetition rate: the mean current (potential drop in electrolyte) decreased by 54-55% when pulse frequency increased from 2 to 30 Hz. Implants performance in terms of the peak amplitudes is summarized in a Supplementary Table 2.

**Table 2.** Mean electric potential during 10 ms pulse in diluted saline (1.56 mS or 640 Ohm*cm) 20µm above the device, generated by various photodiode arrays under 1.64 mW/mm^2^ illumination of the Landolt C pattern with a gap width of 48µm, at pulse frequencies of 2 and 30 Hz.

| Model | Frequency, Hz | 20 μm Pixels | | | 30 μm Pixels | | | 40 μm Pixels | | |
| --- | --- | --- | --- | --- | --- | --- | --- | --- | --- | --- |
|  |  | Bright (mV) | Dark (mV) | Contrast (%) | Bright (mV) | Dark (mV) | Contrast (%) | Bright (mV) | Dark (mV) | Contrast (%) |
| No Shunt | 2 | 40.2 ± 0.1 | 31.3 ± 0.1 | 22 | 42.4 ± 0.1 | 33.8 ± 0.1 | 20 | 48.6 ± 0.1 | 39.5 ± 0.2 | 19 |
| No Shunt | 30 | 18.6 ± 0.1 | 14.6 ± 0.1 | 22 | 19.0 ± 0.1 | 16.0 ± 0.1 | 16 | 22.4 ± 0.1 | 18.7 ± 0.1 | 17 |
| Shunt | 2 | 39.1 ± 0.1 | 28.9 ± 0.1 | 26 | 38.6 ± 0.2 | 30.0 ± 0.1 | 22 | 48.5 ± 0.5 | 32.1 ± 0.2 | 34 |
| Shunt | 30 | 30.7 ± 0.1 | 22.8 ± 0.1 | 26 | 28.1 ± 0.1 | 21.9 ± 0.1 | 22 | 33.4 ± 0.2 | 23.8 ± 0.1 | 29 |

**Figure 8.**
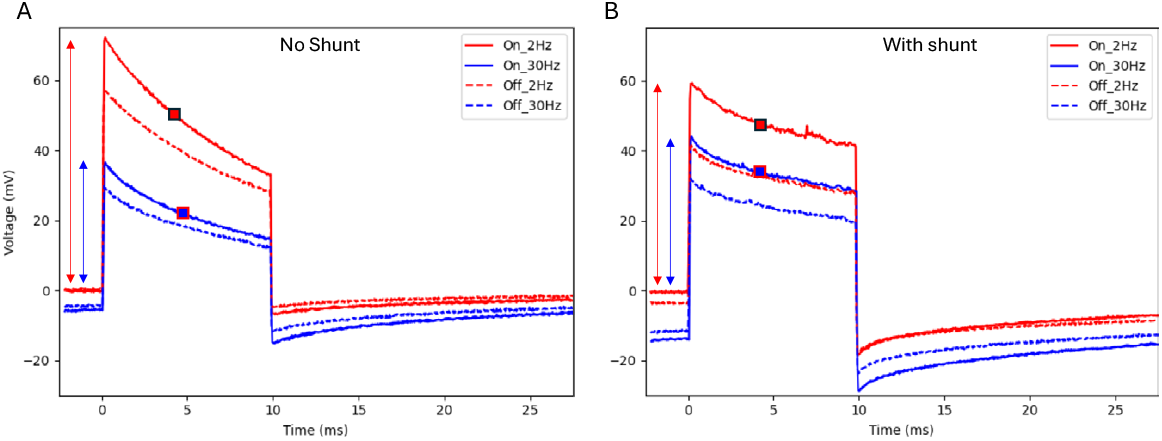
Electric potential measured in a diluted PBS solution (640 Ohm*cm) 20 µm above the implant with 40 m pixels. “On” is above the illuminated part of the letter C and “off” – above the 48 µm gap of the letter. Potential was measured with arrays with and without shunt, using 10 ms pulses at 2 and 30 Hz repetition rate and 1.64 mW/ mm2 irradiance. Amplitude is indicated by the arrows and the mean value during the pulse by a square.

The mean contrast between the gap and the bright part of the letter improved with a shunt from 16-22% to 22-34%, and the contrast level was almost constant during the 10 ms pulses with a shunt. These results demonstrate that incorporating a shunt resistor significantly improved the photodiode’s performance at 30 Hz, better preserved the stimulation strength and increased the contrast of the electric pattern. Further improvement of the contrast, up to about 75%, is expected with pillar electrodes and bipolar pixels (Figure S1B).

## Discussion

As predicted by computational modeling [13,14,19,21], a shunt resistor within each photovoltaic pixel improved the charge injection into electrolyte at high frequency (30 Hz) by discharging the electrodes between the light pulses. It also improved the contrast of the electric patterns by allowing a return current via dark pixels. However, even with a shunt resistor, the contrast with monopolar arrays is still much lower than the nearly 100 % contrast achieved with bipolar pixels of the PRIMA implant [14]. Therefore, the next generation of photovoltaic implants should have return electrodes within each pixel. With a flat geometry, penetration depth of electric field with bipolar pixels decreases with smaller pixel size and becomes insufficient for stimulation of the bipolar cells with pixels below 75 µm [13, 14]. Therefore, the next generation of bipolar arrays will also include pillar electrodes, which penetrate through the subretinal debris layer and reach close proximity to the inner nuclear layer, where bipolar cells reside [20]. Electroplating such pillars on top of the active electrodes can be performed via the shunt resistors, which link to the interconnected return electrode mesh that can be accessed easily through a global wafer connection. The voltage drop across the shunt resistors at the (∼nA) electroplating current is negligible (∼mV).

Fabrication of the MOhm-range resistor on a few µm^2^ area within each pixel requires materials with a layer resistance in the range of MOhm/sq, which translates to resistivity of a few Ohm*cm. Conventional materials used in CMOS fabrication do not cover this range: crystalline silicon is below this range, metal oxides are above it, and doped polysilicon is not well-controlled in this range. Therefore, we had to develop a novel material – doped amorphous silicon (a-Si), which provided resistors in the required range.

Fortunately, fabrication of an a-Si resistor is compatible with the subsequent processes involved in manufacturing of the photodiode arrays, including the alloying and annealing temperature of the metal contacts. In addition, since a-Si is transparent at 880 nm wavelength, this element does not block significant amount of light entering the photosensitive area of the pixel. However, above the a-Si layer, the dielectric coating includes only SiC and not the underlying SiO_2_ layer, and hence its antireflection properties are sub-optimal. Nevertheless, our measurements of the photoresponsivity demonstrated that this small loss did not diminish the photodiode performance – it still remains about 0.53 A/W, similar to the pixels without a shunt resistor (Table 1).

In principle, the optimal value of the shunt resistor depends on retinal resistivity, which is not well known and may vary during retinal degeneration. Fortunately, even though the voltage drop across bipolar cells (stimulation strength) scales linearly with the tissue resistivity, the optimal shunt value (peak of the stimulus strength vs. shunt resistor) changes very little, as shown in the Supplementary Figure 1C. Our current shunt values were optimized for the retinal resistivity of 1000 Ohm*cm [13, 14, 18], but they are not expected to change much even if the tissue resistivity is 4 times higher or lower.

## Acknowledgements

Studies were supported by the National Institutes of Health (R01-EY-035227 and P30- EY-026877), Department of Defense (W81XWH-22-1-0933) and AFOSR (FA9550-20-1-0186). Part of this work was performed at the Stanford Nanofabrication Facility (SNF) and the Stanford Nano Shared Facilities (SNSF), supported by the National Science Foundation under award ECCS-2026822. KM was supported by an RAEng Chair in Emerging Technologies. EB was partially supported by the Rhona Reid Charitable Trust.

## Notes

### Competing Interest Statement

The authors have declared no competing interest.

