## Supplementary material for "Amorphous silicon resistors enable smaller pixels in photovoltaic retinal prosthesis": Figure S1, Figure S2, Table S1, Table S2

### Supplementary Materials

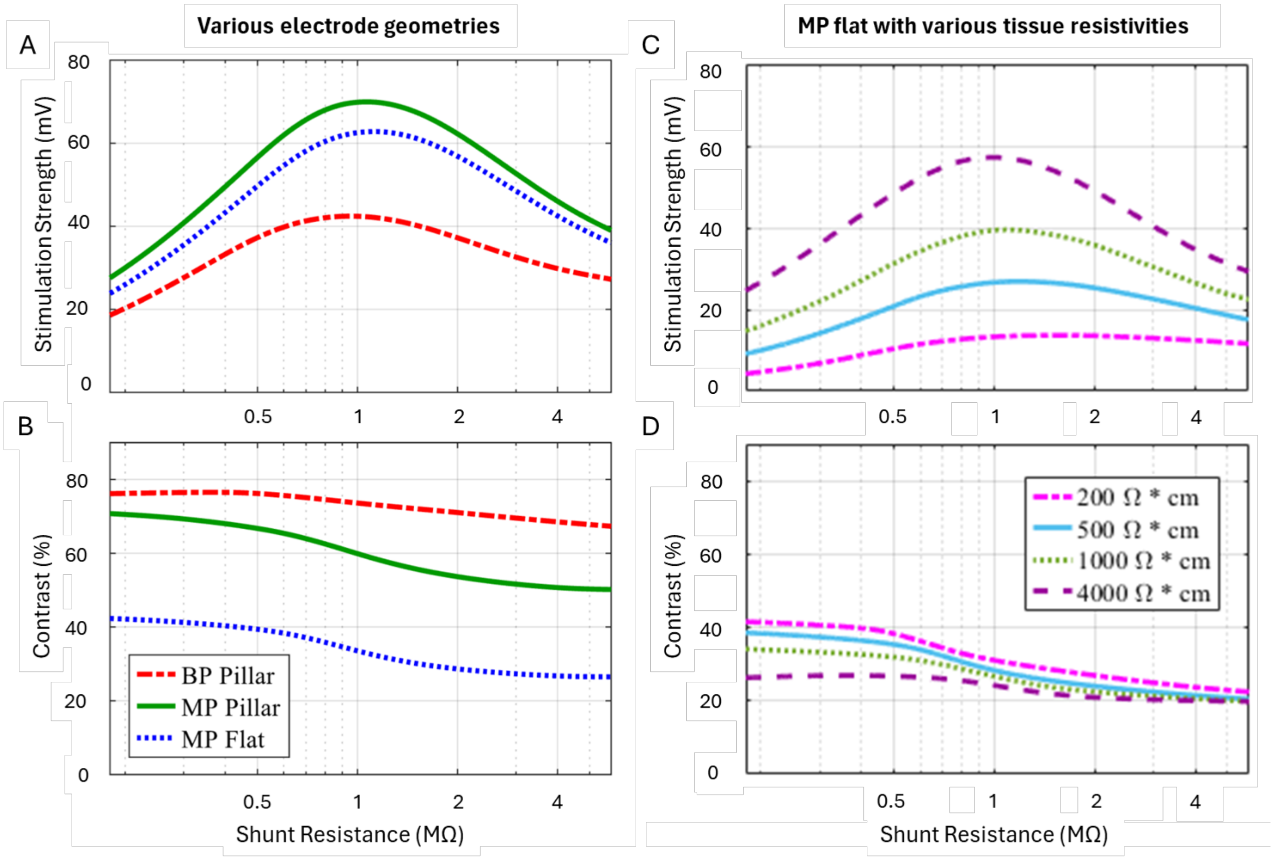


**Figure S1**. Modelled voltage drop and contrast across bipolar cells (35 - 87 μm above the implant) under Landolt C pattern with a gap width of 1.2 pixel size at 1.64 mW/mm$^2$ irradiance and repetition rate of 30 Hz, calculated for 40μm pixels array. (A) Potential drop across bipolar cells as a function of shunt resistance for various electrode configurations: monopolar flat, monopolar pillar and bipolar pillar. (B) Contrast between the gap and bright part of Landolt C. (C) Potential drop across bipolar cells as a function of shunt resistance for flat monopolar pixels, with various retinal resistivities. (D) Contrast between the gap and bright part of Landolt C for conditions shown in C.

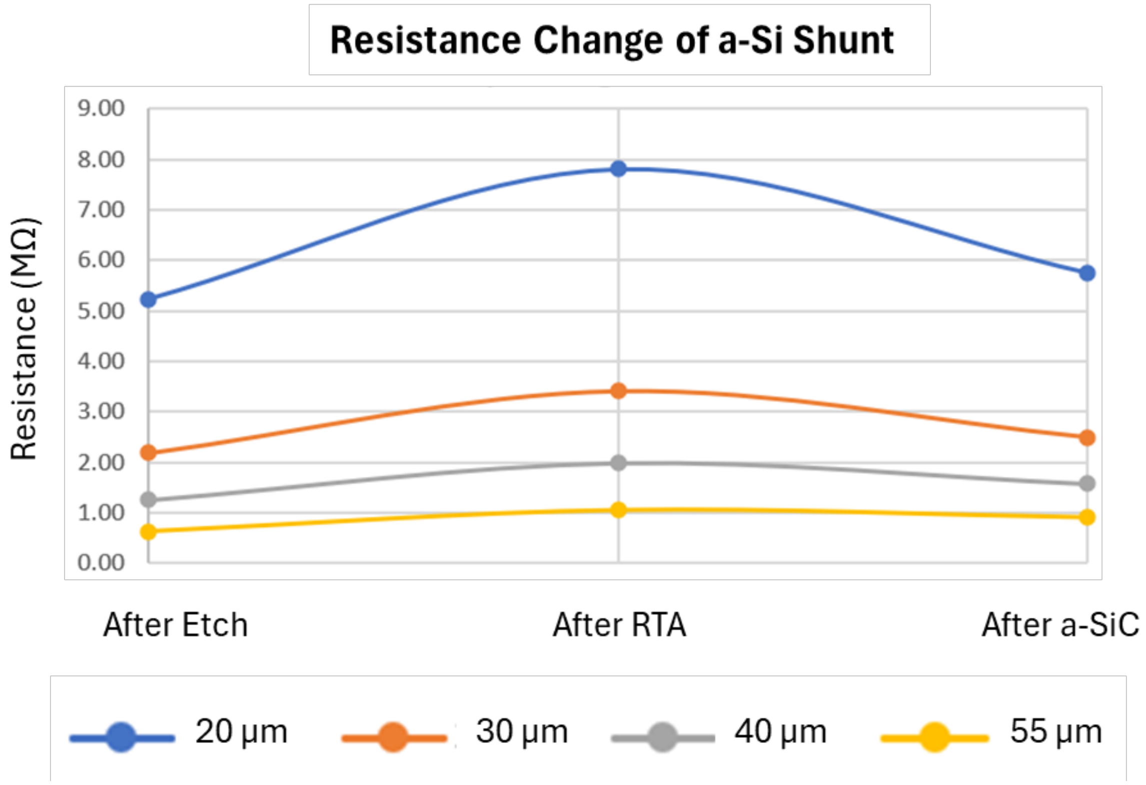


**Figure S2**. Resistance changes of the amorphous silicon layer corresponding to the shunt resistor patterns in the 20, 30, 40, and 55 µm pixels after Rapid Thermal Annealing (RTA) at 425 °C for 30 minutes to improve the ohmic contact between the titanium and amorphous silicon layer, and after adding the a-SiC anti-reflection layer for the insulation purpose.

| Pixel size (μm) | No a-Si LPCVD, no shunt $(\mu m^2)$ | a-Si LPCVD, no shunt $(\mu m^2)$ | a-Si LPCVD, shunt $(\mu m^2)$ |
|---|---|---|---|
| 20 | 164 | 187 | 187 |
| 30 | 413 | 398 | 398 |
| 40 | 778 | 771 | 758 |
| 55 | 1545 | 1533 | 1506 |

**Table S1**. Photosensitive area of various pixels.

| | | *20 μm Pixels* | | | *30 μm Pixels* | | | *40 μm Pixels* | | |
|---|---|---|---|---|---|---|---|---|---|---|
| **Model** | **Frequency, Hz** | **Bright (mV)** | **Dark (mV)** | **Contrast (%)** | **Bright (mV)** | **Dark (mV)** | **Contrast (%)** | **Bright (mV)** | **Dark (mV)** | **Contrast (%)** |
| No Shunt | 2 | **57.9 ± 0.2** | **44.2 ± 0.1** | **24** | **62.6 ± 0.2** | **48.3 ± 0.2** | **23** | **72.1 ± 0.2** | **56.5 ± 0.4** | 22 |
| No Shunt | 30 | **32.3 ± 0.2** | **25.2 ± 0.2** | **22** | **32.1 ± 0.1** | **26.1 ± 0.2** | **19** | **36.5 ± 0.2** | **29.2 ± 0.2** | 20 |
| Shunt | 2 | **48.6 ± 0.2** | **36.3 ± 0.2** | **25** | **50.5 ± 0.3** | **36.9 ± 0.2** | **27** | **60.5 ± 0.5** | **41.5 ± 0.2** | 31 |
| Shunt | 30 | **38.4 ± 0.2** | **28.7 ± 0.1** | **25** | **40.1 ± 0.4** | **28.6 ± 0.1** | **29** | **44.2 ± 0.2** | **32.0 ± 0.2** | 28 |

**Table S2**. Peak electric potential at the beginning of the 10 ms pulse 20μm above the device in diluted saline (1.56 mS, 640 Ohm*cm), generated by various photodiode arrays under 1.64 $mW/mm^2$ illumination of Landolt C pattern with a gap width of 48μm, at pulse frequencies of 2 and 30 Hz.
